# ANNAVP, using neural networks to predict neutralization efficiency of antibodies against viral strains and to cluster strains by protein sequence

**DOI:** 10.1101/2020.09.21.307074

**Authors:** Ghiță Iulian Cristian

## Abstract

Studying viral antibody neutralization data is a complex task and knowledge relating to the effectiveness of a particular antibody to particular strains of viruses cannot easily be extrapolated to other new, related strains. We have developed ANNAVP, a software that uses neural networks to model viral protein data. ANNAVP uses supervised or unsupervised learning and viral protein sequence data to form correlations between different strains and to predict the effectiveness of neutralizing agents against them.

## Introduction

The effectiveness of a treatment against a virus strain can be measured in multiple ways. Among them is the IC_50_ value (µg/mL), representing the antibody concentration in a serum necessary to reduce infectivity by half (Montefiori, 2004). The effectiveness of antibodies depends on many factors, such as their binding to a particular protein forming the viral envelope. The binding affinity between two proteins cannot be directly deduced from their amino acid sequences, due to their secondary and tertiary structures also contributing to said affinity. Difficulties are especially brought by the tertiary structure, whose deduction from the amino acid sequence is an on-going problem in bioinformatics (Ghouzam et al., 2016; Qian & Sejnowski, 1988; Rost & Sander, 1993).

Machine learning algorithms are useful when inferring relationships within large sets of data which lack obvious patterns. One such area of research is determining the effectiveness of antibodies against strains of the HIV-1 virus. Antibodies that confer total immunity against this virus have been difficult to find, due to the binding site of the protein responsible for interaction with the target cell being hidden from antibodies by sugars and by the rest of the protein. The envelope glycoprotein gp12 suffers a conformational change when the virus is near a cell, which exposes the binding site that interacts with the CD4 receptor on the cell surface (Pancera et al., 2010).

Among previous published studies that are using machine learning to predict neutralization data for HIV-1, notable is Buiu et al., 2016, which implemented a feedforward neural network using a Levenberg–Marquardt backpropagation algorithm. With this project, I expanded upon that study to implement other algorithms to achieve the same objective and to also use self-organizing maps to do cluster analysis on viral proteins.

I developed ANNAVP (Artificial Neural Network for the Analysis of Viral Proteins), an offline software with user interface, written in Matlab^™^ (R2012A) using Matlab’s Neural Networks Toolbox (MathWorks^®^, Natick, MA, USA). This software allows the design of multiple types of neural network architectures, using different training algorithms and different ways to codify the amino acid sequences used for training.

## Implementation

The input data used by the software are FASTA files containing the amino acid sequences of the viral protein under study, corresponding to multiple viral strains and CSV files, containing IC_50_ values (or other neutralization quantization values) of antibodies for each strain. The sequences of each FASTA entry can be viewed in the interface with the coordinates of its glycosylation sites marked, where the consensus sequence of the sites is NX(S/T), with X being any amino acid other than proline (An et al., 2009). Before being used by an algorithm, amino acid data must be codified in a machine-readable format. Four types of codifications were implemented: codification A, which converts amino acids in integers from 0 to 25; codification A-6 which consists of six values ranging from 0 to 10 where each value represents an approximation of a physical property (hydrophobicity, volume, charge, aromatic side chain, hydrogen bonds) and a correction bit; codification A-9 is a feature based grouping of amino acids (Brusic et al., 1998); representation B uses six values where each represents the value of a property in its appropriate unit of measurement (volume, bulkiness, flexibility, polarity, aromaticity, charge) (Khudyakov, 2008; Milik et al., 1998). Neutralization data from the CSV files can be viewed as plots of IC_50_ values against each strain or as coverage plots, which show for each serum concentration what percentage of the entries in the file are under that value.

The architecture of the neural network can be set as feedforward or as a self-organizing map. For both types, the number of training steps can be set, which determines how many training attempts the algorithm undertakes.

For amino acid codification type A, the feedforward network type has a number of inputs equal to the viral protein sequence length, a single layer consisting of a user set number of hidden neurons and a target neutralizing value. For the other types of protein codification, the network has two layers and a number of inputs equal to the product between the protein length and the number of values used by the codification type. For each amino acid, each input corresponding to a property is connected to an intermediate network from layer 1, with the number of hidden neurons set by the user. The outputs of all the networks from layer 1 are summed in the network which forms layer 2, whose output is the neutralizing value. The values of the data set can be changed to classes, where each value is converted to one of three classes if it is within certain numerical limits set by the user. The data set as a whole is split up in training, validation and testing subsets. A variety of training algorithms are available and the system’s GPU and CPU multithreading can be used for training.

A self-organizing map is a network which uses unsupervised learning algorithms to organize data according to similarities (Kohonen, 1990). Here, the viral strains are organized in clusters, where each cluster corresponds to a neuron. The number of neurons in the network is set by the user, as is the connection pattern of each neuron with its neighbours (hexagonal, square or random). Furthermore the distance function and initial neighbourhood size can be set.

The resulting trained neural networks are saved in MAT files from where they can be loaded and used. A number of analysis tools are available for each type of network. For feedforward networks, the regression plot for each of the training, validation and testing sets can be plotted. Furthermore, partial least squares (PLS) regression offers plots for the estimated mean square prediction error, for per cent variance against PLS components and a plot for fitted to observed response. Lastly, a sensitivity analysis function allows the running of a selected network with each of its inputs being set to zero one at a time and the difference in performance between each result and the performance at the initial training to be plotted. This allows the identification of key amino acids in the protein structure, essential for matching the target output. A network can be used together with a FASTA file of proteins to generate a CSV file with predicted neutralization values.

For self-organizing maps, one option for analysis is the plotting of SOM Hits, which depicts the network, where each of its neurons is marked with the number of viral strains that have been assigned to it. The View Clusters function generates a table where each column represents a network neuron with its number as the header and its content being the list of strains assigned to it, with the columns ordered by the number of viral strains contained in it. Using the network with a file containing viral proteins, it will generate a file with the list of the strains and the number of the neuron to which each strain was assigned.

## Results

The software was tested by training and comparing the performances of multiple neural networks with different architectures, training algorithms and data codifications. A data set of amino acid sequences pertaining to the variable region of the HIV-1 envelope glycoprotein gp120 from 4907 strains was used to train feedforward networks and self-organizing maps. Furthermore, a set of neutralization data for the NIH45-46 broadly neutralization antibody from the HIV Sequence Database (Kuiken et al., 2003) was used for supervised learning of the feedforward networks (Figure 1).

**Figure 1.**
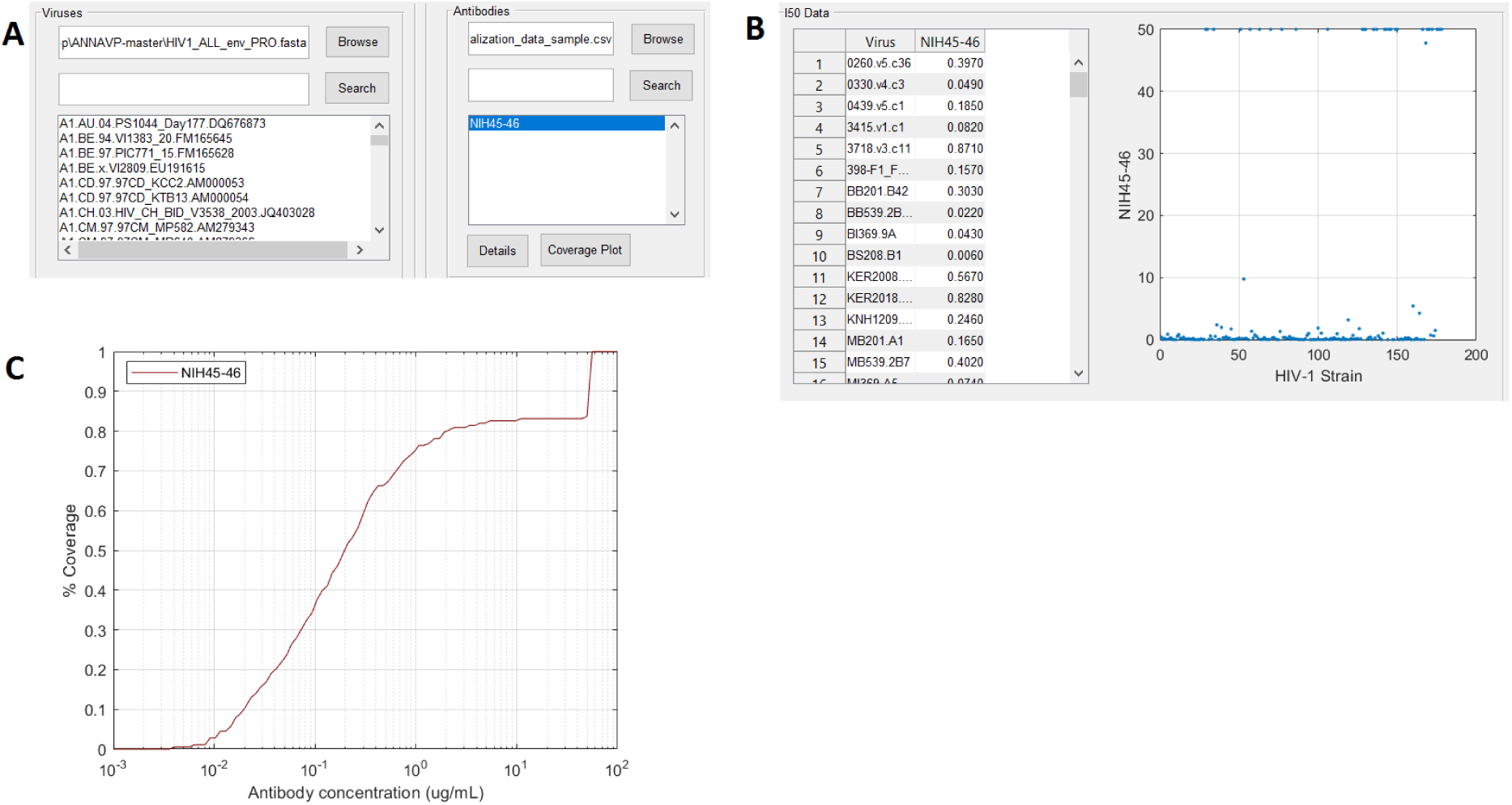
**A** – Loading protein and neutralization data in the UI. **B** – Plot of I_50_ values of NIH45-46 for each viral strain. **C** – Coverage plot of neutralization data shows the percentage of I_50_ values under each concentration from 0.001 µg/ml to 100 µg/ml.

Of the feedforward networks, the best performing one had a layer of 15 hidden neurons, amino acid codification A, the data split as 60% for training, 20% for validation and 20% for training and a Levenberg-Marquardt training algorithm. Network performance was calculated using the mean squared error performance function given by the formula:

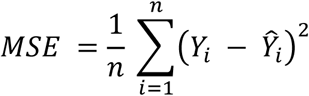

Where *Y* is the vector of target outputs and *Ŷ* is the vector of outputs generated by the network. Performance is best when the *MSE* is the minimum, which was 8.1305 · 10^−28^ for the network mentioned above, as shown in Figure 2. Pearson’s R coefficients for the outputs of the network were 0.82989 for the training set, 0.94285 for validation set and 0.99201 for testing set. Sensitivity analysis revealed that the amino acid coordinates with the most impact on performance were 283, 1071, 1078, 1326 and 1368. Partial least squares regression reduced the data set to incremental numbers components and performed least squares regression on them. For the best performing network, a number of 8 components explained close to 100 per cent of the variance in the data.

**Figure 2.**
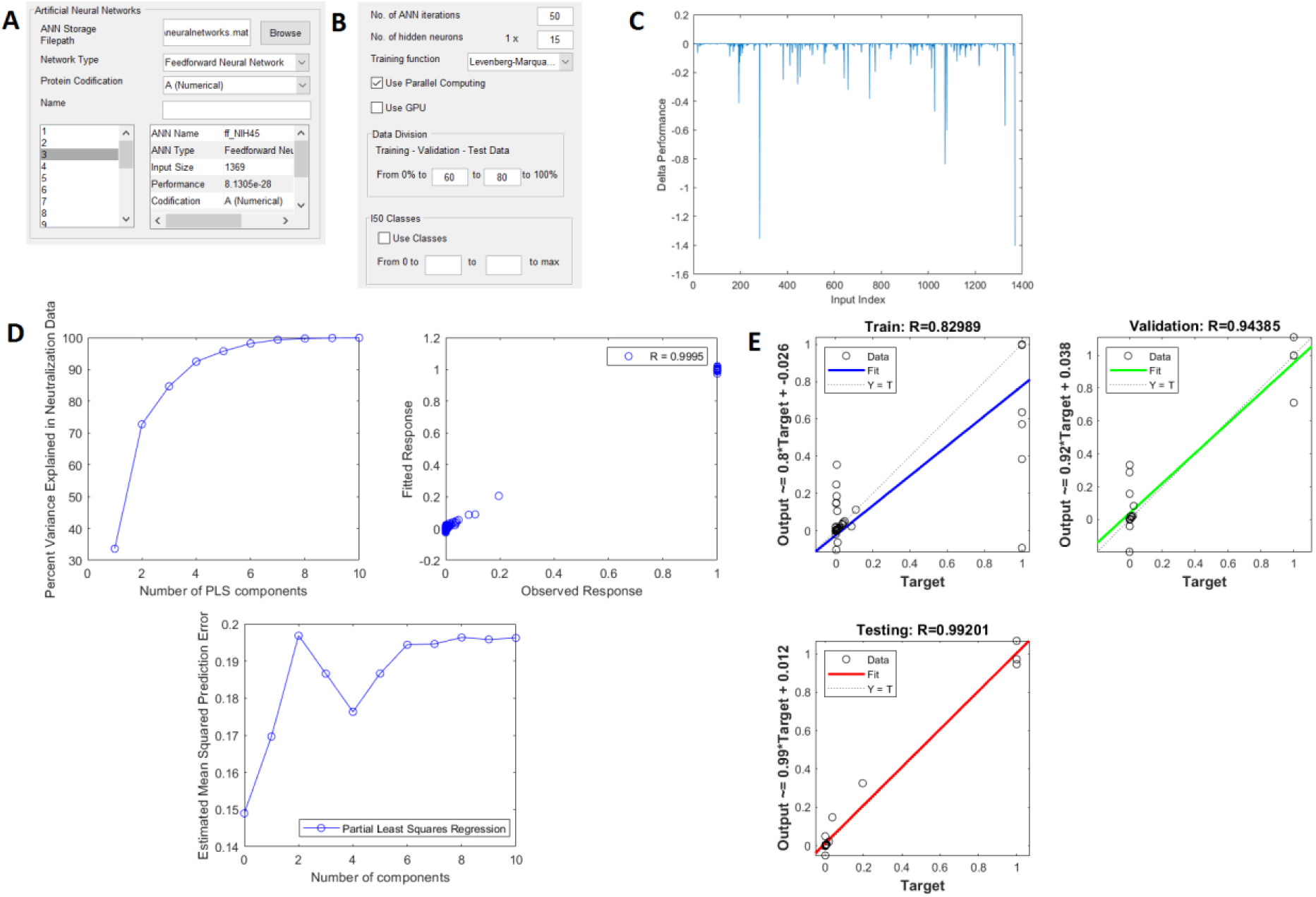
Details for the best performing feedforward neural network. **A** – Parameters for the selection of the network storage file location, type of network, protein codification and details of the networks stored in the file. **B** – Parameters for setting up the feedforward neural network. **C** – Sensitivity analysis; each spike is a network input corresponding to a coordinate in the viral protein which, when set to 0, caused a drop in performance in the network; the length of a spike represents the drop in performance. **D** – PLS analysis; upper left is variance in data as the number of PLS components is increased from1 to 10; it reaches close to total explicative power for at least 8 components; top right is the fitted to observed response for 10 components and Pearson’s R coefficient; bottom is the *MSE* for each number of components. **E** – Plots of target I_50_ values against network output values and R coefficients for each of the three segments of data.

Self-organizing maps cluster the data by assigning each entry in the set to neurons in the network. To asses the performance of each network, multiple alignment with T-coffee was used (Notredame et al., 2000). All the strains assigned to each neuron were aligned and the score of each aligned cluster was compared the score of a sample of the same size, consisting of random strains. The best performance was found for a hexagonal map with 64 neurons, as shown in Figure 3. Data was clustered mainly in 26 neurons where each cluster had more than 15 strains, while the rest of the neurons had at most 4 strains. The alignment scores of the largest 26 clusters were averaged. The average score of each cluster was 97, while the average score for a set of random strains was 87.

**Figure 3.**
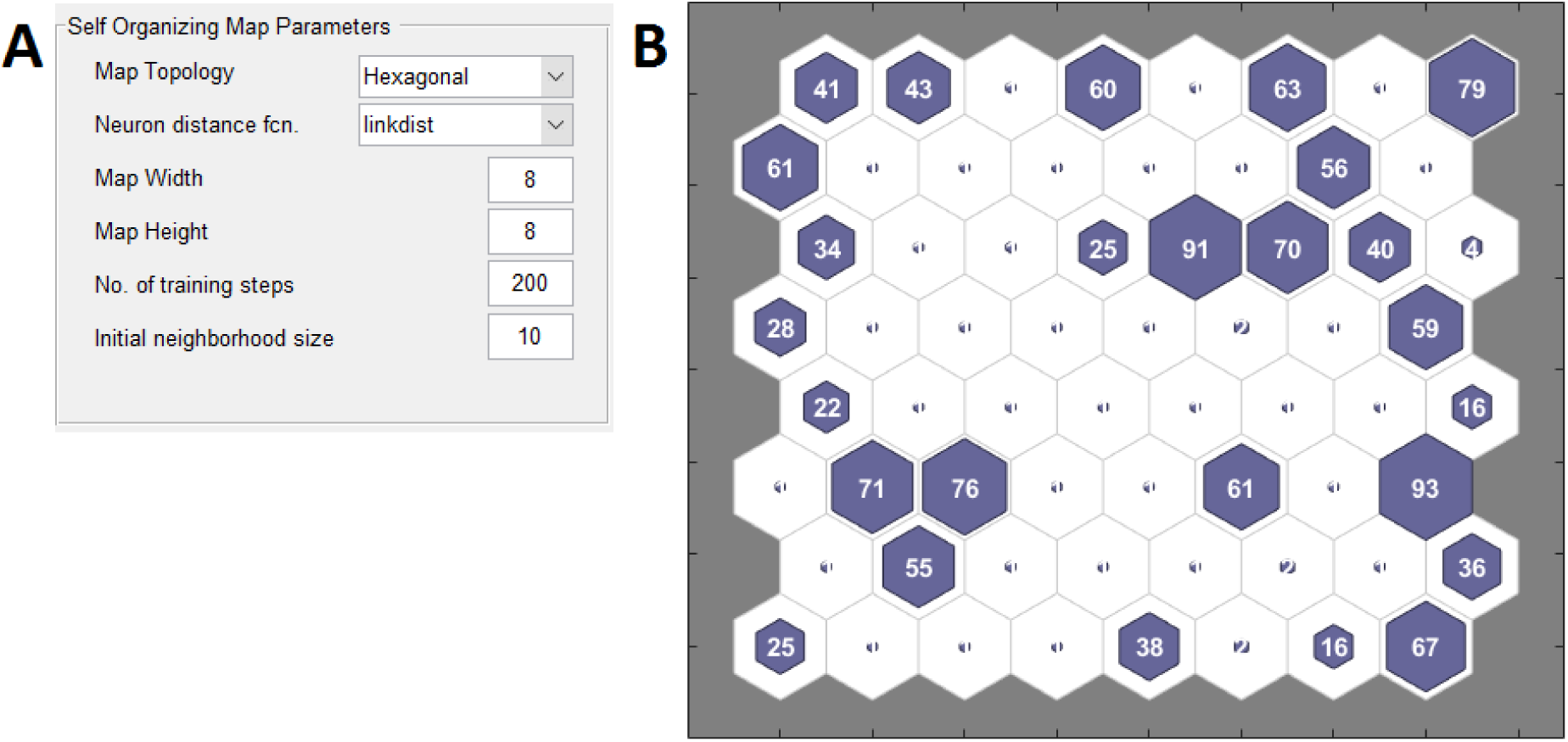
**A** – Parameters used to set the best performing self-organizing map. **B** – Diagram of the self-organizing map; each hexagon is a neuron with the number within indicating the how many viral strains have been assigned to it.

## Discussion

The goal of this paper was to develop interface based software for designing neural networks for the prediction of viral neutralization data and for the clustering of protein sequences from different viral strains. ANNAVP allows the testing of different combinations of network architectures, training algorithms and protein codifications to achieve the best model of the data. The data that can be used is not limited to HIV-1 or to viruses. Proteins from other organisms can be used for modelling; however the software was created with HIV-1 in mind. This software is expected to facilitate the development of more refined and specialized modelling techniques for particular subjects of study.

## Availability

The source code is available on GitHub at https://github.com/icghita/ANNAVP.

## Acknowledgements

I acknowledge the contributions of Prof. Dr. Cătălin Buiu of the Politechnic University of Bucharest, Faculty of Automation and Computers, for providing guidance and the data for the development of this project.

